# EVALUATION OF STAINING FOR FIBRONECTIN ISOFORMS IN LIVER DISEASES

**DOI:** 10.1101/2025.05.14.653452

**Authors:** Stefan Hamelmann, Juan C. Sarmiento, Thekla Bung, Mariam Menhem, Ari J Cohen, Gretchen Galliano, Paul Thevenot, Alan Burshell, Inaam A. Nakchbandi

**Author notes:** Shared first authors (Stefan Hamelmann, Juan C. Sarmiento). Funding Sources:* German Research Council (DFG: NA400/5; NA400/7; NA400/9; NA400/10); Max-Planck Society (M.KF.A.BIOC0001). Correspondence:* Inaam Nakchbandi, MD, FACP, Institute of Immunology, University of Heidelberg, Im Neuenheimer Feld 305, 2. OG, 69120 Heidelberg, Germany.

## Abstract

Fibronectin is an ubiquitous extracellular matrix protein, that is produced by the hepatocytes contributing to circulating fibronectin, and by the hepatic stellate cells contributing to fibrotic bundles. Fibronectin is required for the accumulation of collagen type I, the main fibrotic collagen in liver disease. It is presumed that cellular fibronectins including EDA-containing, EDB-containing and oncofetal fibronectin are the ones localized to the fibrotic bundles, since these isoforms are increased in the presence of fibrosis and cirrhosis in patients with liver disease. The aim of this work was therefore to evaluate the distribution of the various isoforms and determine whether they are detected in the hepatocytes too or whether they are indeed limited in their expression to fibrotic bundles.

Total fibronectin was detected in hepatocytes particularly in healthy liver tissue. In advanced disease, fibronectin staining seemed diminished. All three isoforms, EDA-, EDB-, and oFN were detected in hepatocytes or the fibrotic bundles, with EDB being the most abundant. Interestingly, the isoforms did not fully account for all of fibronectin. Staining patterns confirmed that the three isoforms are not limited to the fibrotic bundles, but are also found in the hepatocytes. This contradicts the common notion that cellular fibronectins precipitate and are localized in the matrix, and explains why they can be measured in the circulation. Finally, there is some convergence between the presence of the isoforms in the fibrotic bundles and the elevation of their levels in the blood stream at least in chronic hepatitis C and cholestatic liver disease. This is in line with a high degree of complexity of fibronectin distribution and effects.

## Introduction

Fibronectin is an extracellular matrix glycoprotein produced by many mammalian cell types ^1^. Its importance can be implied by early embryonal death in knockout mice due to failure of normal vascular development ^2^. It is required in health and can affect disease development making it an interesting molecule to study.

### Structure and isoforms

Fibronectin is a glycoprotein consisting of two peptide chains attached to each other via cysteine bridges ^3^. Each chain consists of a series of modules, type I, II and III. Three type III modules are of particular interest. The extra-domain A (ED-A, also called EIII-A as an allusion to the fact that it is a type III domain) and the extra-domain B (ED-B or EIII-B) can be present and give their name to the fibronectin isoform (EDA-fibronectin contains an ED-A domain, and EDB-fibronectin contains EDB) ^4^. There is little distinction in the literature between these two isoforms and they are mostly combined under the term cellular fibronectin, which however can also refer to fibronectin that includes both modules: ED-A and ED-B. It is thought that most of fibronectin in the matrix between the cells belongs to this group ^5, 6^. Both modules are absent in the dominant circulating form of fibronectin that is produced by the hepatocytes and secreted into the circulation called plasma fibronectin ^7, 8^. From the bloodstream, this isoform can infiltrate various organs including the kidney and bones ^9, 10^. This isoform also is able to contribute fibronectin to cancer ^11, 12^.

The third type III module of interest is the so called variable region, because it can be present in different lengths ^13^. This variable region needs to be present in at least one of the two chains for fibronectin to be secreted ^14^. Fibronectin can undergo various post-translational modifications, one of which is the presence of an O-glycosylation in the amino-terminal part of the variable region ^15^. A specific antibody detects this O-glycosylation and defines the so called oncofetal fibronectin, named so because it is expressed prenatally in the placenta in large amounts ^16, 17^. Some also use the name oncofetal fibronectin for the isoform EDB-fibronectin ^18^. We will however use the presence of O-glycosylation as the prerequisite for the isoform to be called oncofetal fibronectin (oFN).

While some scientists continue to differentiate between “plasma” (for the circulating form) and “cellular” fibronectin (for the matrix form), this distinction has been increasingly called into question, because we have shown that isoforms viewed as cellular fibronectins can be detected in the bloodstream ^6, 15, 19^.

### Functions

Fibronectin stimulates proliferation, inhibits apoptosis and supports differentiation ^4, 12^. It is able to do so by binding to a large number of different receptors including integrins, toll-like receptors and the hyaluron receptor ^1^. In addition, its ability to form fibrils is required for collagen type I accumulation ^20^. Thus inhibiting fibronectin fibril formation suppresses collagen amount in the matrix ^21, 22^. Many functions of fibronectin have been described. It is involved in different physiologic processes such as angiogenesis and blood clot formation ^12^. It modulates the response to some bacterial infections, can affect bone health, and supports tumor progression ^4, 9, 10, 12, 22^.

### Role in health and disease

Circulating fibronectin consists almost exclusively of plasma fibronectin and originates from the hepatocytes where it is produced in parallel to albumin and therefore reflects the ability of the liver to synthesize proteins ^23^. It is this fibronectin that can infiltrate the bone contributing to its integrity and that is detected in the glomeruli too ^9, 10^. In liver disease, the hepatic stellate cells become activated. These myofibroblasts then produce the extracellular matrix that consists of fibronectin and collagen. Too much of this matrix: i.e., fibrosis, leads to distortion of the normal structure of the basic functional units called the hepatic lobule. As matrix accumulates, the blood is no longer able to flow unhinged between the portal vein carrying various nutritional and toxic molecules and the inferior vena cava of the systemic circulation. This interferes with the ability of the liver to filter the blood and replenish various components ^24, 25^. Initial evidence *in vitro* that hepatic stellate cells, once stimulated, produce mainly the fibronectin isoforms: EDA-, or EDB-containing fibronectin and oncofetal fibronectin (oFN) was confirmed by their increase in the circulation, possibly escaping from the matrix, in various liver diseases ^15, 26^. Indeed, one isoform (oFN) was found to be increased in the circulation of patients with cholestatic liver disease and to mediate hepatic disease-associated osteoporosis by diminishing osteoblast function ^15, 27^.

Increased levels of all three cellular forms of fibronectin in the circulation in patient with chronic hepatitis C also showed a correlation with the grade of fibrosis, with the highest values particularly of EDA-fibronectin and oFN detected in liver cirrhosis ^19^. The aim of the current work is therefore to evaluate liver sections for staining patterns of the various cellular isoforms, try to identify particular staining pattern associated with the various liver diseases and attempt to draw conclusions about the origins of the isoforms.

## Methods

### Staining Protocols and Immunohistochemistry

Liver tissue samples were obtained from the pathology department at Ochsner Health in New Orleans, LA. These were collected as part of usual tissue collection for adequate patient care and embedded in paraffin. The use of these samples for research purposes was based on approval by the institutional review board of the Ochsner Clinic Foundation (Approval nr. 2017/608). Sections (5 μm thick) were cut with a rotation microtome (Leica RM2145), mounted on SuperFrost Plus slides (#03-0600, R. Langenbrinck GmbH) and stored at 4°C until staining. For deparaffinization and rehydration, the sections were submerged in xylol three times, each submersion lasting for 3 minutes, washed in descending ethanol series, each for 3 minutes (3 x 100 %,1 x 95% and 1 x 80%), followed by washing with ddH^2^O for 5 minutes. Quenching was performed using 100 mM Ammonium chloride and 100 mM glycine for 10 minutes. This was followed by washing the slides in phosphate-buffered saline (PBS) for 2 minutes. For antigen retrieval, sections were heated in sodium-citrate-buffer (2.94 g/l tri-sodium citrate-dihydrate, 0.5 ml/l Tween 20, pH 6.0) in a water bath for 3 h, left to cool down to room temperature for 20 minutes, and washed twice with PBS-T (PBS+ 0.5 ml/l Tween 20, pH 7.0) for 2 minutes. Immunofluorescence staining for plasma fibronectin was performed by using a total fibronectin antibody #MFCD00162291 (clone F3648, Sigma-Aldrich) applied at a dilution of 1:100 for 2 h. Staining for the different isoforms was performed by using the following antibodies all left on the samples overnight: EDA-fibronectin #MFCD00164562 (clone FN-3E2, Sigma-Aldrich, dilution 1:50); EDB-fibronectin #S-FN12 (clone C6, Sirius Biotech, dilution 1:25); oncofetal fibronectin (oFN) #ATCC-HB-9018 FHCR-1-2813/FDC-6, dilution 1:25).

After washing twice with PBS for 2 minutes, the sections were incubated with secondary antibodies and DAPI (0,01 mg/ml) for 1 h at room temperature. Total fibronectin was detected using Cy5 goat anti-rabbit Star635P (#ST635P-1002-500UG, Abberior, dilution 1:100); EDA-fibronectin was detected using Cy5 goat anti-mouse Star635P (#ST635P-1001-500UG, Abberior, dilution 1:100); EDB-fibronectin was detected using Cy5 goat anti-mouse Star635P (#ST635P-1001-500UG, Abberior, dilution 1:100) and oncofetal fibronectin using Cy5 goat anti-mouse Star635P (#ST635P-1001-500UG, Abberior, dilution 1:100). After washing twice with PBS for 2 minutes, sections were dehydrated by immersion for 5 minutes 95 % Ethanol repeated three times and 5 minutes 100 % ethanol repeated three times too. The sections were covered with Mowiol (Dissolve 5 g Mowiol (4-88 reagent, Calbiochem) in 20 ml PBS, add 10 ml glycerine, heat at 50°C for 30 minutes in a water bath, centrifuge at 5000 g for 30 minutes followed by storage at -20°C). Sections were then stored at 4°C. Stained slides were analyzed and photographed using a fluorescence microscope ECLIPSE Ti (Nikon) and the attached software NIS-Elements (version 4.50, Nikon). Preparation of the images was performed using the program FIJI (Wayne Rasband, NIH; ImageJ).

#### Ethics Statement and Patient Consent

The study was approved by the Ochsner institutional review board under number 2017/608 and consisted of evaluation of liver samples with subject identification codes already assigned and for which consent was obtained as part of a prior approved study.

## Results

### Characteristics of the group

A total of 39 patients was evaluated. Six patients with chronic hepatitis C (Hep C), one patient with chronic hepatitis B, 11 with hepatocellular carcinoma (HCC), 5 patients with non-alcoholic steatohepatitis (NASH), 2 primary biliary cirrhosis (PBC), 5 primary sclerosing cholangitis (PSC), 5 chronic liver transplant rejection, 4 metastatic adenocarcinoma, and 1 hemangioma patient. Because changes in chronic hepatitis B could not be confirmed in at least two more patients, we excluded the patient from the analysis.

### Total fibronectin

Total fibronectin was detected throughout the sections. The findings will be presented for the hepatocytes and for the fibrils for each disease entity.

The first panel represents a section from a patient with chronic hepatitis C without primary antibody addition. This sample therefore serves as a negative control to exclude any non-specific binding by the secondary antibody. In the absence of a primary antibody, the laser intensity during microscopy can be set such as to correctly detect weak signals in the samples exposed to both the primary and the secondary antibody (Figure 1A).

**Figure 1.**
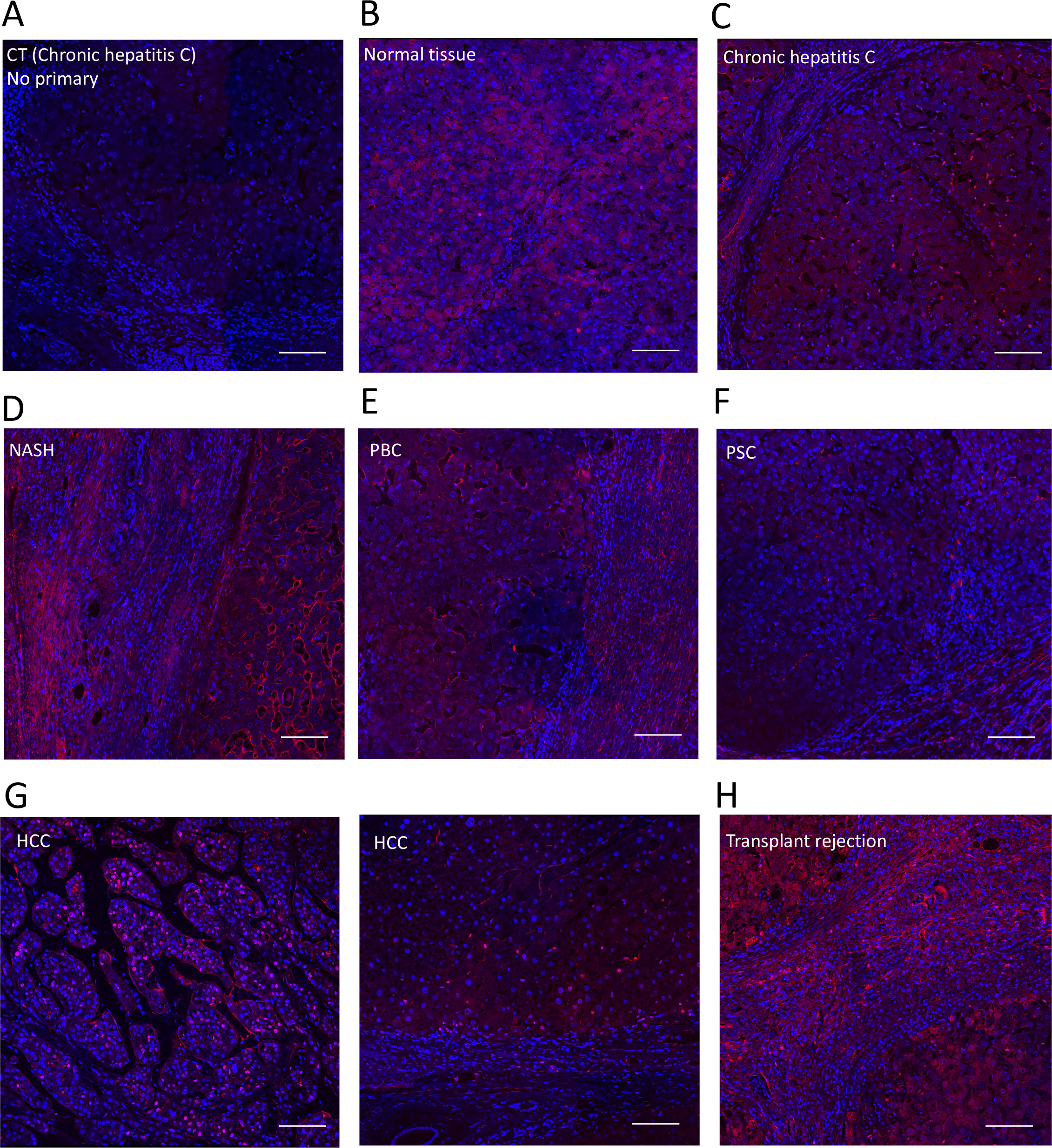
Total fibronectin staining and distribution in various liver diseases. A. Non-specific staining by the secondary antibody was excluded in sections of a patient afflicted with chronic hepatitis C. Shown is an area that includes hepatocytes and a fibrotic bundle. Nuclei are stained with DAPI. B. An area is shown from the patients with hemangioma. Normal liver tissue is seen with diffuse fibronectin staining (in red) in hepatocytes. C. In chronic hepatitis C, hepatocytes are stained and fibrils are seen on the left. Between the hepatocytes, the sinusoids are visible and their margins are stained for fibronectin. D. In non-alcoholic steatohepatitis, strong and diffuse staining in the fibrils is seen, the sinusoids are well visualized giving the impression of less staining in the hepatocytes. E. and F. Diffuse staining in PBC and decreased hepatocyte staining in PSC, but the bundles are well-stained in both disease entities. G. An area of hepatocellular carcinoma is shown. the margins and central areas in the cells are stained, but in the adjacent tissue without tumor, changes similar to C (chronic hepatitis C) are seen. H. In transplant rejection, the fibrotic bundles are very cellular with marked staining for fibronectin. Bars represent 100 μm.

Adenocarcinoma and hemangioma were not detected in the sections. These samples therefore can serve as liver-healthy controls, particularly the hemangioma sample. The staining was homogeneous and diffuse in the cytoplasm of hepatocytes (Figure 1B), and no fibrils were seen in the absence of fibrotic disease.

In chronic hepatitis C, intracellular fibronectin staining was homogeneously distributed in the hepatocytes. The fibrotic bands contained dispersed fibronectin-stained fibril-like structures. A network was visible that sometimes bordered the hepatocytes consistent with sinusoids as shown in figure 1C.

In NASH there were more nuclei in the fibrotic bundles consistent with slightly more prominent inflammation compared to chronic hepatitis C (Figure 1D).

In PBC and PSC, fibronectin fibrils were visible in the fibrotic bundles, in particular in the vicinity of bile ducts, with associated hypercellularity consistent with active inflammation (Figure 1E and F).

In contrast, hepatocellular carcinoma (HCC) showed heterogeneity between the various samples and sometimes within the same sample. Occasionally, single cells or groups of cells showed enhanced staining compared to the remaining hepatocytes (Figure 1G).

In transplant rejection, the hypercellularity in the fibrotic bundles was more pronounced than in chronic hepatitis C and seemed to be diffuse compared to cholestatic liver diseases, in particular PSC. Staining was pronounced (Figure 1H).

In summary, fibronectin is homogeneously present throughout the cytoplasm of most hepatocytes. It is fibrillar in the presence of fibrotic bundles, and seems increased in HCC in groups of hepatocytes and in the fibrotic bundles.

### EDA-containing fibronectin

In comparison to total fibronectin staining, there is a larger degree of variability between the various diseases.

In the normal tissue surrounding adenocarcinoma and angioma, the hepatocytes seemed diffusely stained with some heterogeneity between the hepatocytes within a lobule. In general, the distribution of staining in the lobule seemed similar to total fibronectin, albeit more granular in the cells (Figure 2A+B).

**Figure 2.**
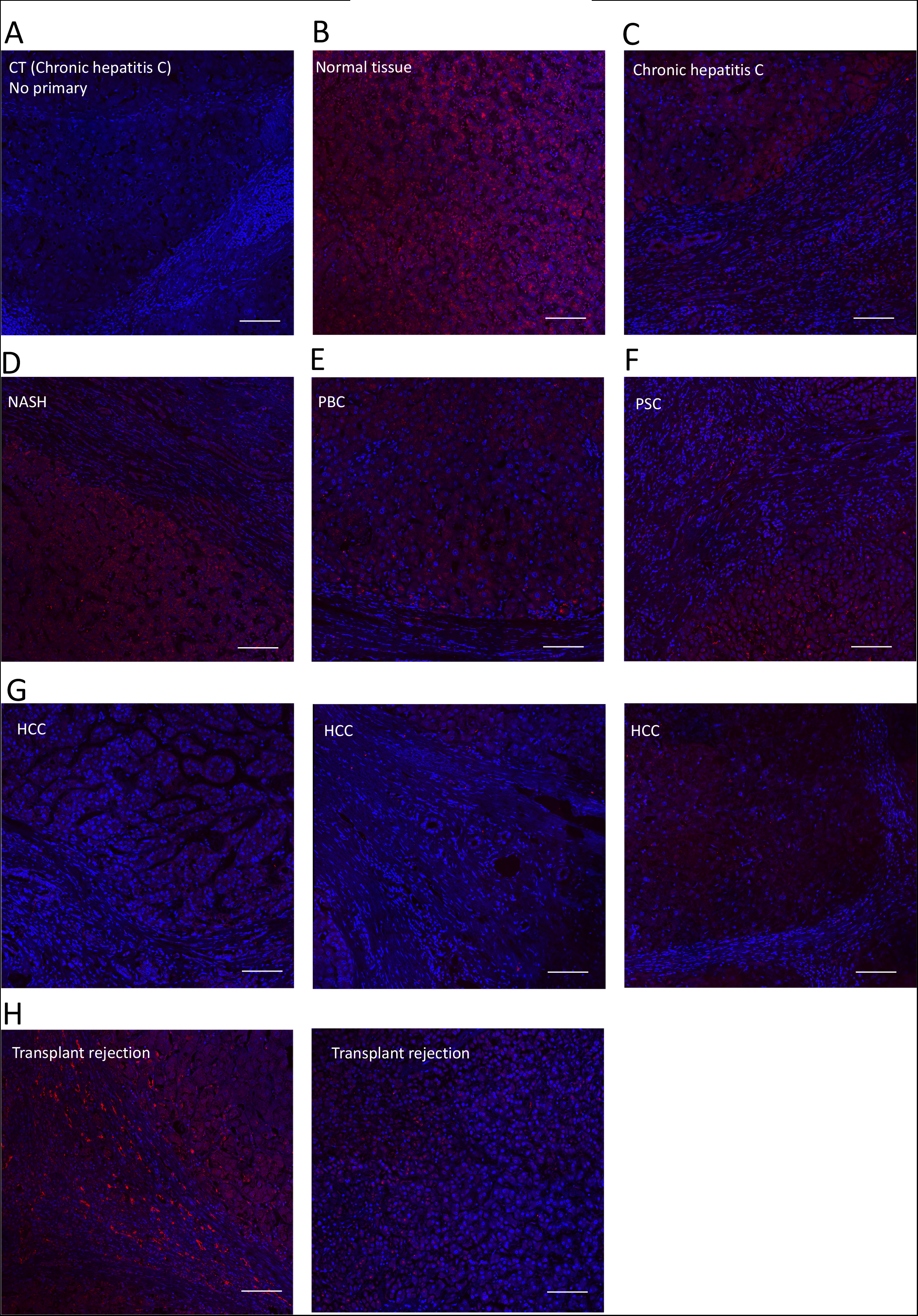
EDA-fibronectin staining and distribution in various liver diseases. A. Non-specific staining by the secondary antibody was excluded in sections of a patient afflicted with chronic hepatitis C. Shown is an area that includes hepatocytes and a fibrotic bundle. Nuclei are stained with DAPI. B. An area is shown from the patients with hemangioma. Normal liver tissue is seen with diffuse fibronectin staining (in red) in hepatocytes. C. In chronic hepatitis C, hepatocytes are stained and fibrils are seen on the right. Some staining is seen in the bundles. D. In non-alcoholic steatohepatitis (NASH), strong and diffuse staining in the fibrils is seen. E. and F. Diffuse staining in PBC and PSC with occasional clumps in the hepatocytes. The bundles show some staining. G. An area of hepatocellular carcinoma is shown. Little staining is detected. H. In transplant rejection the fibrotic bundles are highly cellular with clear staining for EDA-fibronectin in the bundles. Bars represents 100 μm.

In chronic hepatitis C, punctate staining sometimes close to the cell membrane was detected in the hepatocytes. Furthermore, unlike total fibronectin, fibrils seemed unstained but the cytoplasm of cells of the biliary ducts were stained as were rare cells in the fibrils, presumably fibroblasts suggesting that some fibroblasts contain EDA (Figure 2C).

In NASH, the staining was always punctate and diffuse in the cytoplasm of almost all hepatocytes (Figure 2D). Only rarely was staining detected in the fibrils, but when present then it is clear as shown in the figure. A similar picture was seen in PBC and PSC (Figure 2E+F).

In HCC, the staining seemed mostly cytoplasmic in the hepatocytes and occasionally punctate. The fibrils seemed prominent and long, with little staining (Figure 2G).

In transplant rejection, the staining was mostly cytoplasmic, but clumps were seen too. The fibrils were well delineated and EDA was detected where a lot of fibrils were present (Figure 2H).

In summary, EDA shows mostly a granular pattern in the hepatocytes. It is also found in the fibrils, but the amount is much less than total fibronectin, suggesting it contributes little to the fibrils except in transplant rejection.

### EDB-fibronectin

In normal liver tissue adjacent to hemangioma and adenocarcinoma, inhomogeneous punctate staining was seen in some but not all hepatocytes, and appeared sometimes close to the nucleus (Figure 3A+B).

**Figure 3.**
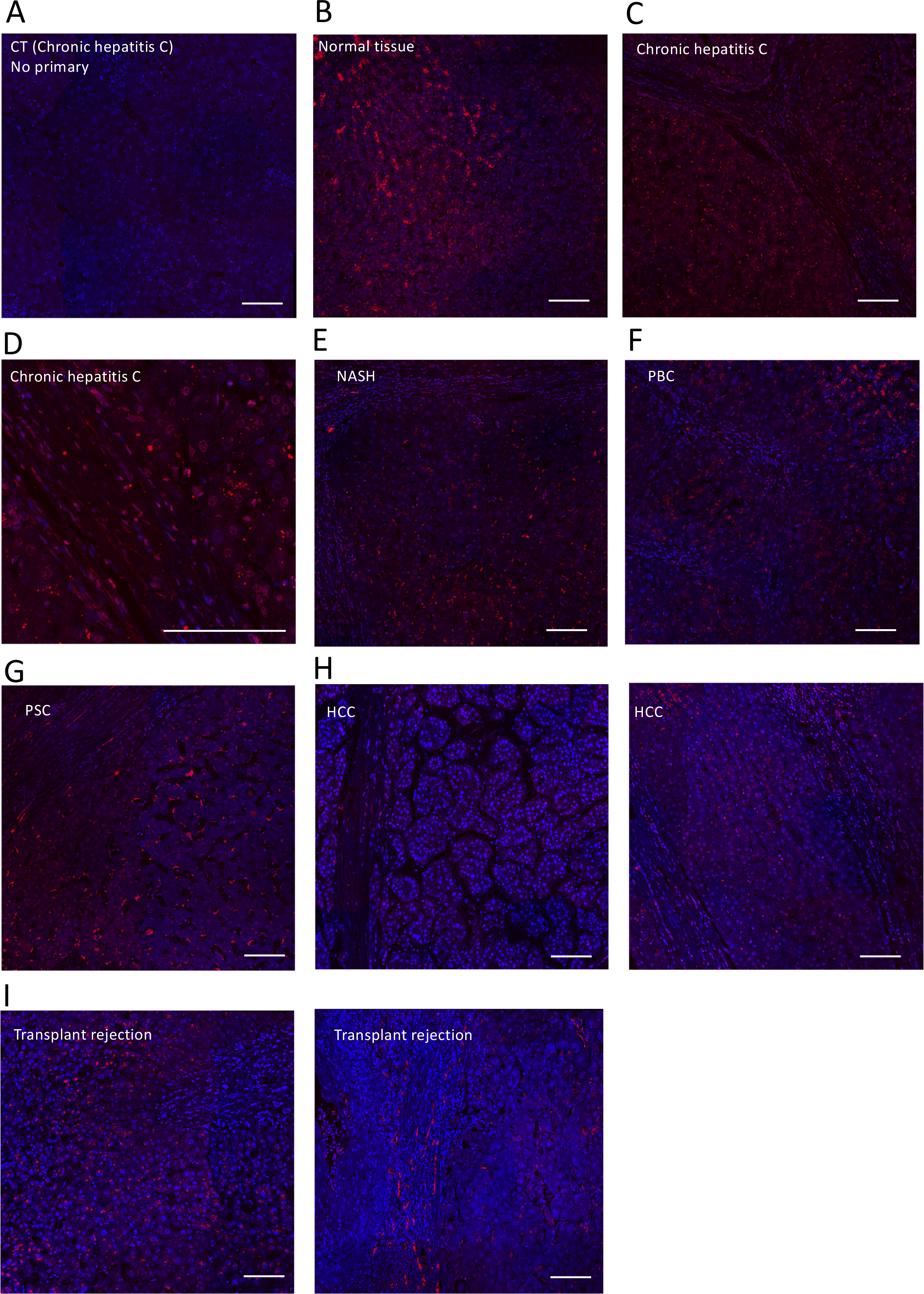
EDB-fibronectin staining and distribution in various liver diseases. A. Non-specific staining by the secondary antibody was excluded in sections of a patient afflicted with chronic hepatitis C. Shown is an area that includes hepatocytes. Nuclei are stained with DAPI. B. An area is shown from the patient with hemangioma. Normal liver tissue is seen with granular fibronectin staining (in red) in hepatocytes. C. In chronic hepatitis C, hepatocytes were stained and fibrils are seen in the middle. Some staining is seen in the bundles. D. A smaller area is enlarged to show the staining in the fibrotic bundles, some of which is associated with nuclei, i.e., in fibroblasts. E. In non-alcoholic steatohepatitis (NASH), strong, diffuse and occasionally clumpy staining in the fibrils is seen. F. and G. Granular staining in PBC and clumpy staining in PSC is seen in hepatocytes. The bundles show particularly strong and fibrillar staining in PSC, and not in PBC. H. An area of hepatocellular carcinoma is shown. Little staining is detected. In the non-tumor adjacent tissue on the right panel, the fibrotic bundles showed very short fibrils or perinuclear staining of fibroblasts. I. In transplant rejection the fibrotic bundles are very cellular with fibrillar staining for EDB-fibronectin in the bundles. Bars represent 100 μm.

In chronic hepatitis C, there seemed to be less staining in the hepatocytes, but if present it was punctate and not homogeneously distributed. In the fibrotic bundles, perinuclear staining was noted suggesting that a cell type present in the bundles expresses EDB, but some fibrils were also stained (Figure 3C+D).

In NASH, the staining was mostly inhomogeneous and punctate in the hepatocytes, with some stained and some unstained, and some areas showing more staining than others. In fibrils, occasional cellular/perinuclear staining is detected in addition to staining in the fibrils (Figure 3E). In PBC and PSC, the hepatocytes were inhomogeneously stained and seemed granular. While only very short fibrils were seen in the fibrotic bundles in PBC, longer fibrils were detected in PSC (Figure 3F+G).

In HCC, very little homogeneous perinuclear staining was seen. The EDB-stained fibrils seemed occasionally long (Figure H+I).

In transplant rejection, staining was either dispersed or close to the nucleus and perinuclear staining was detected in the fibrotic bundles with fibrils of variable lengths (Figure J+K).

In summary, EDB-fibronectin resulted in punctate staining in hepatocytes that was occasionally perinuclear. Fibrotic bundles also showed staining that was both perinuclear or with distinct fibrils seen. This staining was more pronounced than EDA, but still less than total fibronectin.

### oFN-fibronectin

Little oFN was detected in normal tissue. If present, staining was homogeneous within each cell, cytoplasmic and rarely perinuclear. The picture shown is of one of these areas (Figure 4A+B). Similarly, in chronic hepatitis C, minimal staining in the hepatocytes was detected but the fibrotic bundles showed distinct peri- and paranuclear staining suggesting staining in fibroblasts (better seen at higher magnification) (Figure 4C+D).

**Figure 4.**
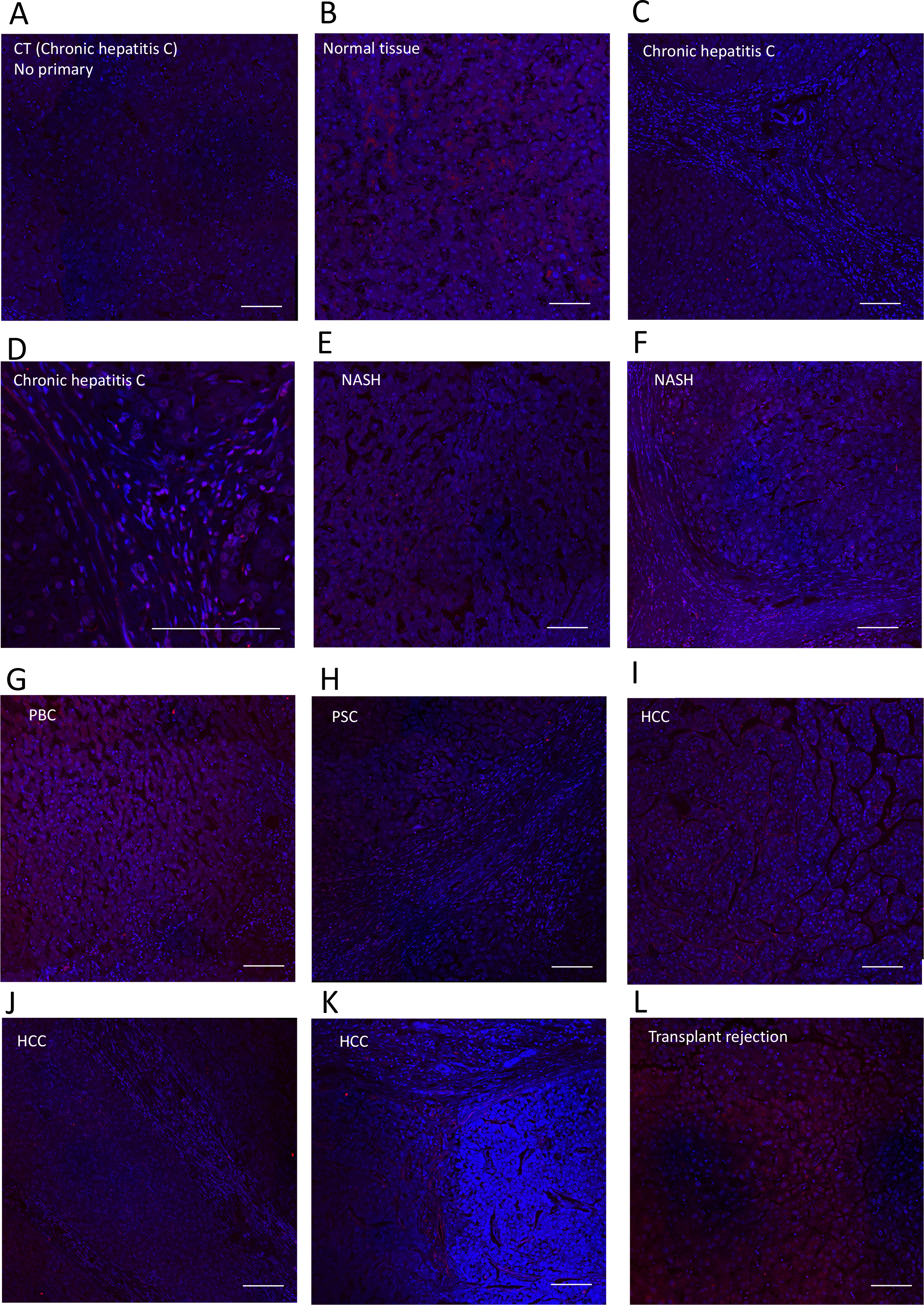
Oncofetal fibronectin (oFN) staining and distribution in various liver diseases. A. Non-specific staining by the secondary antibody was excluded in sections of a patient afflicted with chronic hepatitis C. Shown is an area that includes hepatocytes. Nuclei are stained with DAPI. B. An area is shown from the patient with hemangioma. Normal liver tissue is seen with granular punctate fibronectin staining (in red) in hepatocytes, but not all hepatocytes show the same degree of staining. C. In chronic hepatitis C, little staining in the hepatocytes but more in the fibrils can be detected. The picture with higher magnification better shows the staining in the fibrotic bundles, E-F. In non-alcoholic steatohepatitis (NASH), little staining in hepatocytes but stronger staining in the fibrils is seen. G-H. Diffuse staining in PBC and almost no staining in PSC in the hepatocytes. Instead, some staining is detected in the bundles in PSC. I. An area of hepatocellular carcinoma is shown. Little staining is seen. J+K. In the non-tumor adjacent tissue of HCC, the fibrotic bundles seem strongly stained with short fibrils and perinuclear staining of fibroblasts. L. In transplant rejection, the hepatocytes show diffuse staining for oFN seemingly similar to PBC. Bars represent 100 μm.

In NASH, groups of hepatocytes showed weak, patchy, sometimes granular staining occasionally localized close to the nuclei on one side consistent with staining of oFN in the Golgi apparatus, where the glycosylation of fibronectin that defines oFN takes place. Fibrils are also seen (Figure 4E+F). In PBC, hepatocyte staining was diffuse, while in PSC fewer hepatocytes were stained. Fibrils in PSC showed short stained segments and fibroblastic staining (Figure 4G+H).

In HCC, staining was very heterogeneous with perinuclear staining on one side in some hepatocytes but the fibrils were relatively well-stained (Figure 4I+J+K).

In transplant rejection, there was variable staining that affects clusters of hepatocytes but not all of them (Figure 4L).

In summary, oFN is occasionally cytoplasmic and sometimes localized on one side of the nucleus. It is detected in fibrotic bundles particularly in chronic hepatitis C and HCC.

## Discussion

Total fibronectin and three domains that characterize isoforms were stained in common liver diseases. This setup helped define the distribution of the isoforms in comparison to total fibronectin in health and in various liver pathologies. It should be noted however that fibronectin actions are complex. On one hand, genetic decrease in fibronectin enhanced experimental liver fibrosis, by increasing TGF-β availability. On the other hand, inhibiting fibronectin fibril formation pharmacologically and consequently decreasing the attachment and accumulation of collagen I, reduced collagen I amounts in the liver and suppressed fibrosis progression ^21, 25^. Thus, no direct conclusions can be drawn based on staining intensity alone.

In healthy liver tissue, total fibronectin was diffusely distributed in the cytoplasm of many hepatocyte. In contrast, both EDA- and EDB-fibronectin were punctate and more limited in their distribution with the staining mostly localized close to the nucleus. This suggests that the isoforms first accumulate in the endoplasmic reticulum and are either released very quickly out of the cell or that the amounts in the cytoplasm are lower than required to be detected. oFN was barely detectable. Even though total fibronectin was diffusely cytoplasmic, some hepatocytes and their margins showed more staining than others, irrespective of their locations in normal liver tissue. This suggests that not all hepatocytes secrete similar amounts of fibronectin. Hepatocytes produce circulating fibronectin, which diminishes in parallel with albumin production ^23^. A reduction in total fibronectin production therefore reflects a deterioration in protein synthesis by the hepatocytes in various diseases, and explains the weaker staining in chronic hepatitis C or loss of staining seen in PSC (Figure 1C and 1F).

Considerably more fibrils were stained for total fibronectin than the isoforms. This suggests that the isoforms contribute some, but not all of the fibrillar fibronectin. Furthermore, fibronectin detected in the fibrils is most likely produced by the fibroblasts located in the fibrils (as can be discerned in Figure 3D or 3G), but could also originate from the hepatocytes. In contrast to EDA which diminished in hepatocytes in liver disease (except in NASH, Figure 2D, which looks similar to healthy tissue, Figure 2B), it was noted that EDB remained clearly expressed in the hepatocytes in all diseases evaluated and was detected in some, but not all fibrils in a distribution and form similar to total fibronectin, albeit less pronounced with some fibrillar bundles that were not stained. A considerable amount of fibrillar fibronectin contained EDB in HCC and in chronic hepatitis C, in which the prominence of EDB-stained fibrils is in line with the increase in circulating EDB-fibronectin reported in the latter patients ^19^. Thus, EDB is part of the fibrils in some diseases despite the lack of change in EDB mRNA reported by others ^28^.

Little oFn expression was found in healthy liver tissue in hepatocytes in line with the detection of circulating levels of oFN in these subjects ^15^. In the absence of any fibrotic bundles, this suggests that isoforms in the blood stream can originate from the hepatocytes. oFN seemed to be expressed in hepatocytes in PBC, which would explain its increase in this disease ^13, 15^. A similar degree of staining in hepatocytes of transplant rejection patients suggests that it could also be elevated in this disease entity. However, all three isoforms increased in the bloodstream in patients with cholestatic liver diseases and chronic hepatitis C ^15, 19^. Since both EDA and EDB were sometimes detected in perinuclear stains in the fibrils in chronic hepatitis C (Figure 2C and 3D), it seems reasonable to suggest that some the increase in the levels of these isoforms in the circulation originates from the fibroblastic cells located in the fibrillar bundles in these patients.

At least in the case of chronic hepatitis C and PSC, the increase of oFN in the circulation is not associated with pronounced staining in the hepatocytes, raising the possibility that in these cases, the fibroblasts in the fibrils contribute to the increase in circulating oFN, and supporting the conclusion reached for EDA and EDB in chronic hepatitis C ^15, 19^. Thus, it is no longer surprising that elevated levels of EDA and oFN most reliably reflected how advanced fibrosis was ^15, 19^. In contrast, in PBC, the major contributor is probably the hepatocyte, where homogeneous staining is detected. Indeed, in this disease, elevated oFN levels were associated with more complications ^15^. Thus, circulating oFN can be released from the hepatocytes, the fibrils, or both. However, the speed of release of fibronectin from the cells could differ in the different diseases preventing a final judgement on which cells contribute in the various diseases. Since transplant recipients show staining in hepatocytes as seen in PBC, it might be feasible to use circulating levels of oFN in this disease entity to define the extent of injury.

In conclusion, all isoforms are expressed in hepatocytes, whereby EDB expression is detected in more cells than either EDA or oFN. All isoforms are also expressed in elongated cells in the fibrils that could be fibroblasts suggesting that these cells also produce them. Comparisons of the expression in hepatocytes and fibrils with findings in the circulation suggest that both cell types, hepatocytes and fibroblasts in the bundles contribute to circulating levels of the isoforms to various degrees depending on the disease. The exact source be it hepatocytes or fibroblasts: i.e., hepatic stellate cells cannot be fully discerned, however, because the staining is not always a reflection of the elevation in the bloodstream. Finally, while EDB fibronectin contributes markedly to fibronectin in the fibrotic bundles, EDA and oFN contribute to a lesser extent.

## Authors contributions

AB, IAN: Conceptualization; SH, JCS, MM, AC, GG, PT, TB: Data curation and/or formal analysis; IAN: Funding acquisition; SH, JCS, TB: Investigation; SH, JCS, TB: Methodology; AB, IAN: Project administration; AB, IAN: Resources; AB, IAN: Supervision; TB: Validation; SH, JCS, TB: Visualization; IAN: Writing - original draft; AB, IAN: Writing - review & editing.

## Acknowledgment

We acknowledge the Biospecimen & Core Research Laboratory of the Ochsner Clinic Foundation. Funding sources: The German Research Council (Deutsche Forschungsgemeinschaft: DFG; NA400/5-1; NA400/5-2; NA400/9); Max-Planck Society (M.KF.A.BIOC0001/K440).

## Declaration of competing interest

The authors declare that they have no known competing financial interests or personal relationships that could have appeared to influence the work reported in this paper.

## Notes

No conflicts of interest exist for any of the authors.

### Competing Interest Statement

The authors have declared no competing interest.

